# Detection of allele-specific expression in spatial transcriptomics with spASE

**DOI:** 10.1101/2021.12.01.470861

**Authors:** Luli S. Zou, Tongtong Zhao, Dylan M. Cable, Evan Murray, Martin J. Aryee, Fei Chen, Rafael A. Irizarry

## Abstract

Allele-specific expression (ASE), or the preferential expression of one allele, can be observed in transcriptomics data from early development throughout the lifespan. However, the prevalence of spatial and cell type-specific ASE variation remains unclear. Spatial transcriptomics technologies permit the study of spatial ASE patterns genome-wide at near-single-cell resolution. However, the data are highly sparse, and confounding between cell type and spatial location present further statistical challenges. Here, we introduce spASE (https://github.com/lulizou/spase), a computational framework for detecting spatial patterns in ASE within and across cell types from spatial transcriptomics data. To tackle the challenge presented by the low signal to noise ratio due to the sparsity of the data, we implement a spatial smoothing approach that greatly improves statistical power. We generated Slide-seqV2 data from the mouse hippocampus and detected ASE in X-chromosome genes, both within and across cell type, validating our ability to recover known ASE patterns. We demonstrate that our method can also identify cell type-specific effects, which we find can explain the majority of the spatial signal for autosomal genes. The findings facilitated by our method provide new insight into the uncharacterized landscape of spatial and cell type-specific ASE in the mouse hippocampus.

## Introduction

In diploid organisms, allele-specific expression (ASE) refers to the imbalanced expression of the two parental alleles for a given gene. ASE has been well-studied in the context of epigenetic phenomena such as genomic imprinting and X-chromosome inactivation (XCI) [1, 2, 3], where expression from one allele is silenced. Spatial patterns of ASE have long been observed as a consequence of XCI in female organisms, where the random silencing of either the maternal or paternal X-chromosome in early development is passed to daughter cells, resulting in visible clusters of ASE [4, 5, 6]. By contrast, although studies in bulk and single-cell RNA-sequencing data have revealed widespread variability in ASE throughout the autosome across tissues and cell types [7, 8, 9, 10, 11, 12, 13, 14, 15, 16, 17, 18, 19, 20], relatively little is known about the prevalence of spatial ASE therein.

Spatial transcriptomics technologies now provide the opportunity to study spatial ASE patterns genome-wide. For example, Slide-seqV2 [21, 22] has high resolution which enables near-single-cell quantification of ASE with 2D spatial information. However, these data are limited by highly sparse read counts in comparison to bulk or single-cell sequencing technologies, which is further exacerbated by the requirement that reads align uniquely to one allele. In addition, cell type, which drives the majority of variability observed in single-cell data, is highly correlated with spatial location, especially in solid tissue [23]. Therefore, it is important to distinguish between spatial and cell type-specific ASE, which could arise from and contribute to distinct underlying biological mechanisms.

Several statistical and computational methods have been developed for studying ASE in bulk and single-cell RNA-seq data [24, 25, 26, 27, 28, 29, 30, 31]. Some focus on estimating allele-specific transcriptional bursting kinetics for individual genes in homogeneous populations of cells [15, 30, 31]. Here, we instead focus on the problem of estimation and inference for the maternal allele probability *p* for a given gene across 2D space, and we consider how p may vary with cell type. To model p in bulk and single-cell RNA-seq, multiple methods have used a beta-binomial framework, which can flexibly account for overdispersion from unknown technical and biological variability [26, 27, 28]. An additional advantage of this model is that it can be parameterized as a generalized linear model (GLM) [32, 33], allowing for maximum likelihood estimation of p while incorporating covariates of interest such as cell type.

The issue of estimating smooth functions from sparsely sampled data has been well-studied [34, 35, 36, 37, 38], and multiple solutions have been developed and implemented as computational methods [39, 40]. In the case of allele-specific spatial transcriptomics data, although the read count measured at individual spatial coordinates may be low, smoothing spline methods can increase power by leveraging information from local neighborhoods of pixels. Generalized additive models are GLMs that incorporate smoothing splines into a regression framework, enabling estimation of the smooth spatial function as well as hypothesis testing for spatial functions deviating from a constant [38, 40].

Here, we present spASE, a computational framework for detecting genes with significant ASE patterns in spatial transcriptomics data. We employ a hierarchical beta-binomial smoothing approach based on thin plate regression splines [36, 41] to estimate 2D allele probability functions and detect spatially significant genes. Given the high correlation between cell type and spatial location in solid tissue, our method permits control for cell type effects as well as any other potential covariates of interest. Through simulations, we confirm the power and false positive rate control of our method even in highly sparse settings such as those observed in allele-resolved spatial transcriptomics. Additionally, we generate allele-specific Slide-seqV2 data from the hippocampus of an F1 hybrid mouse and find that we are able to recover known patterns of ASE due to XCI in highly-expressed X-chromosome genes, both within and across cell types. We further show that our method can detect cell type-specific ASE, which we find can explain most of the spatial signal observed in autosomal genes such as *Ptgds*. Overall, we report new insights into the uncharacterized landscape of spatial and cell type-specific ASE in the mouse hippocampus, thus demonstrating the utility of spASE for detecting known and novel patterns of ASE in spatial transcriptomics.

## Results

### A beta-binomial framework for modeling allele-specific spatial transcriptomics

A statistical challenge for allele-specific spatial transcriptomics is that spatial and cell type effects can be confounded (Figure 1a,b). We therefore developed a computational framework that can account for both these sources of variability. Specifically, we developed a beta-binomial GLM that provides a flexible approach to estimation, inference, and visualization of ASE in spatial transcriptomics. We denoted the counts from the maternal allele for gene *g* and pixel *i* with *Y_gi_* and assumed it followed the distribution:

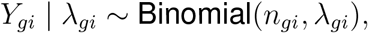

with *n_gi_* the observed total UMI count for gene *g* and pixel *i*, summing both alleles, and λ*_gi_* the probability that a transcript from gene *g* is from the maternal allele. We assumed that λ*_gi_* follows a beta distribution with mean *p_gi_* and variance *ϕ_g_p_gi_*(1–*p_gi_*). Here, *p_gi_* is the mean maternal allele probability and *ϕ_g_* is a gene-specific overdispersion parameter ranging from 0 to 1 that accounts for biological and technical variability not explained by binomial sampling.

**Figure 1:**
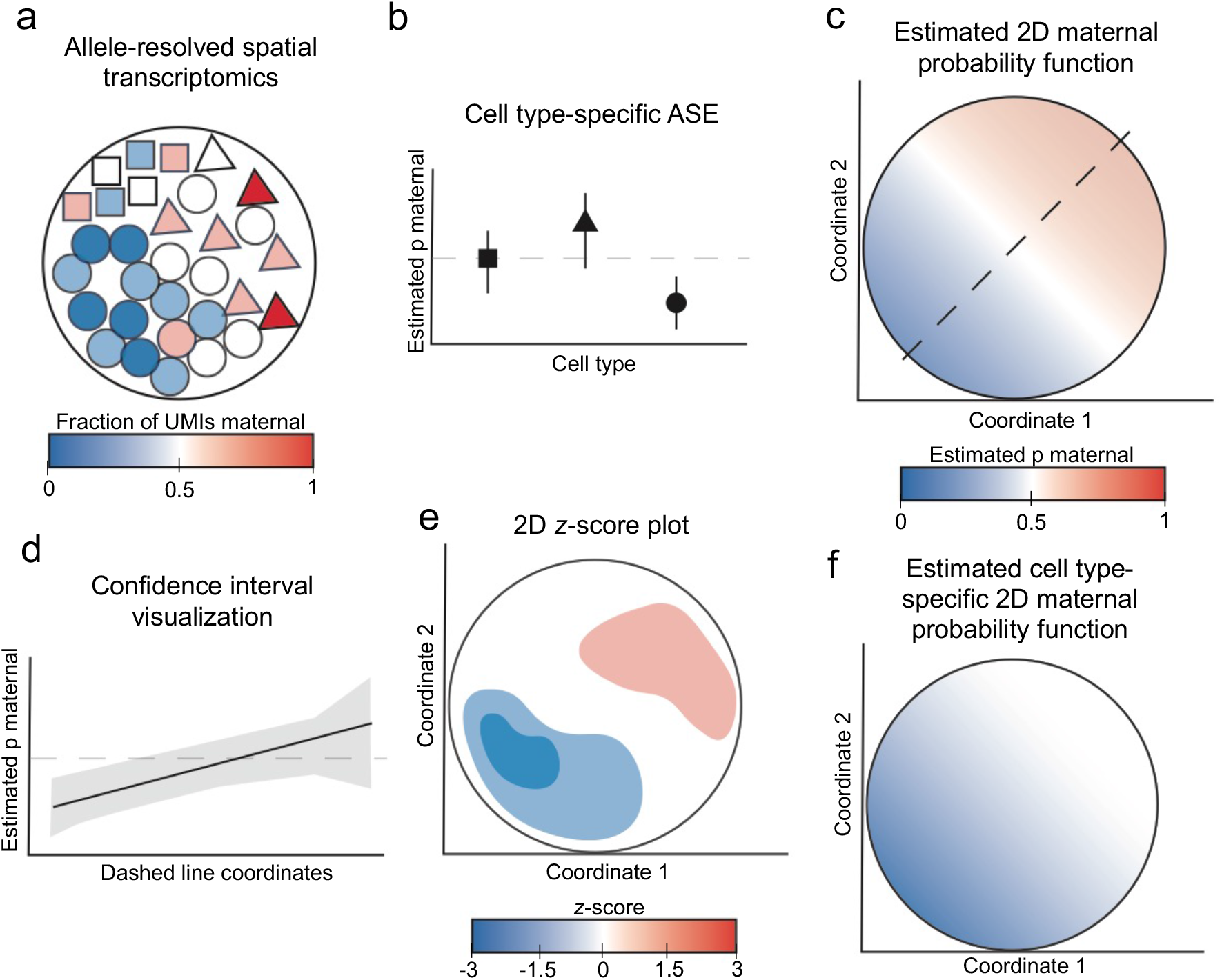
Schematic of detecting allele-specific expression in spatial transcriptomics using spASE. **(a)** Input is allele- and cell type-resolved spatial transcriptomics with UMIs. Each shape represents a different cell type, and the color indicates the fraction of observed UMIs that were from the maternal allele. **(b)** Point estimates and confidence intervals for the estimated maternal allele probability (estimated p maternal) for each cell type. **(c)** Visualization of the estimated maternal probability function, not controlling for cell type. **(d)** Visualization of confidence intervals (gray shaded region) around the MLE in a 1D cross-section. The solid line indicates the estimated maternal probability along the black dashed line from c. Light gray dashed line indicates the null of *p* = 0.5. **(e)** 2D z-score plot visualizing region-level significance of the estimated function from c. **(f)** Estimated cell type-specific function for the circle cell type from a.

To account for spatial and cell type effects, we created a logit-linear GLM,

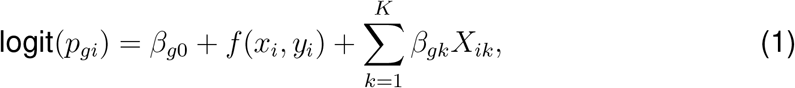

with *x_i_* and *y_i_* the spatial location of pixel *i*, *f*(*x_i_, y_i_*) a smooth function of location, the *X_ik_*’s indicator functions equal to 1 if pixel *i* is from cell type *k*, and the *β_gk_* parameters representing gene-specific cell type effects. Note that the *β_gk_* can be interpreted as the change in log-odds, compared to the reference cell type, of a maternal allele transcript in gene *g* and cell type *k*.

The spatial effect function was modeled as a thin plate spline [36] defined by

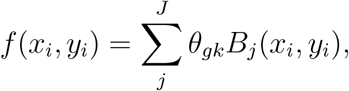

with *B_j_*(*x, y*) the smooth basis function for the spline and *θ_gk_* the gene-specific parameters that define gene-specific spatial effects. With this definition of *f*, all the terms in (1) are linear and define a GLM. We can therefore obtain maximum likelihood estimates (MLEs), standard errors, and confidence intervals for all parameters using GLM theory and software. Furthermore, we can test for spatial effects by performing a likelihood ratio test comparing the model with *f* to a model without space (see Methods for details).

In addition to fitting spatial ASE across all cell types, we can also fit a cell type-specific version of model (1) as

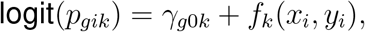

for all pixels *i* belonging to cell type *k*, where *p* and *f* have been modified to depend on cell type. Note that only certain genes within certain cell types provide enough counts and therefore power to fit our cell type-specific spatial model.

After estimating parameters by maximum likelihood for each gene, we can visualize the smooth maternal probability function across 2D coordinates (Figure 1c), and for any given spatial cross-section, we can additionally visualize confidence intervals (Figure 1d). Region-specific significance can also be assessed using 2D *z*-score maps (Figure 1e). Running spASE on individual cell types generates spatial ASE maps for cell type-specific estimation (Figure 1f). spASE uses the likelihood ratio to rank genes according to spatial effects variability.

To evaluate the performance of our method, we generated simulated spatial transcriptomics data under a wide variety of sparsity and overdispersion conditions (Supplementary Figure S1, Methods). We calculated the power and false positive rate, and we computed p-values to detect significant spatial ASE. We observed that power decreased as overdispersion increased (Supplementary Figure S1a); however, we found that power is at least 70% even for genes with high overdispersion (*ϕ* = 0.8) and as few as 50 pixels with low UMI coverage (e.g. less than 10 UMIs per pixel). With at least 100 pixels for a given gene, the power across all scenarios was at least 85%, even with as low as 1 UMI per pixel. Using a p-value threshold of *p* ≤ 0.01, we found that the false positive rate approached the nominal rate of 0.01 as the number of pixels increased, in concordance with the expected asymptotic guarantees of our model (Supplementary Figure S1b,c). We also evaluated confidence interval coverage as a function of sample size and number of UMIs per pixel (Supplementary Figures S2, S3), and we found that the beta-binomial model maintained near-95% coverage across all scenarios.

### spASE identifies spatially-significant ASE genes and smooths over sparse allele-specific spatial transcriptomics signal

To test spASE on allele-specific spatial transcriptomics data, we generated Slide-seqV2 data of an F1 hybrid CAST/EiJ x 129S1/SvImJ (CAST x 129) mouse hippocampus and surrounding region (see Methods). We aligned 150bp reads to a pooled CASTx129 transcriptome and only considered reads that uniquely aligned to one allele. We used RCTD [23] to call cell types using a single-cell RNA-sequencing reference of the mouse hippocampus [42], and we filtered to pixels with a high likelihood of sourcing UMIs from a single cell type (Figure 2a). Based on results from our simulations (Supplementary Figure S1), we filtered genes with non-zero UMI counts on at least 100 pixels. Using these filtering criteria resulted in 4,140 genes for downstream analysis, which were expressed on a median of 210 pixels (IQR: 140-384 pixels) with a median number of UMIs per pixel of 1.06 (IQR: 1.04-1.1 UMIs/pixel) (Supplementary Figure S4).

**Figure 2:**
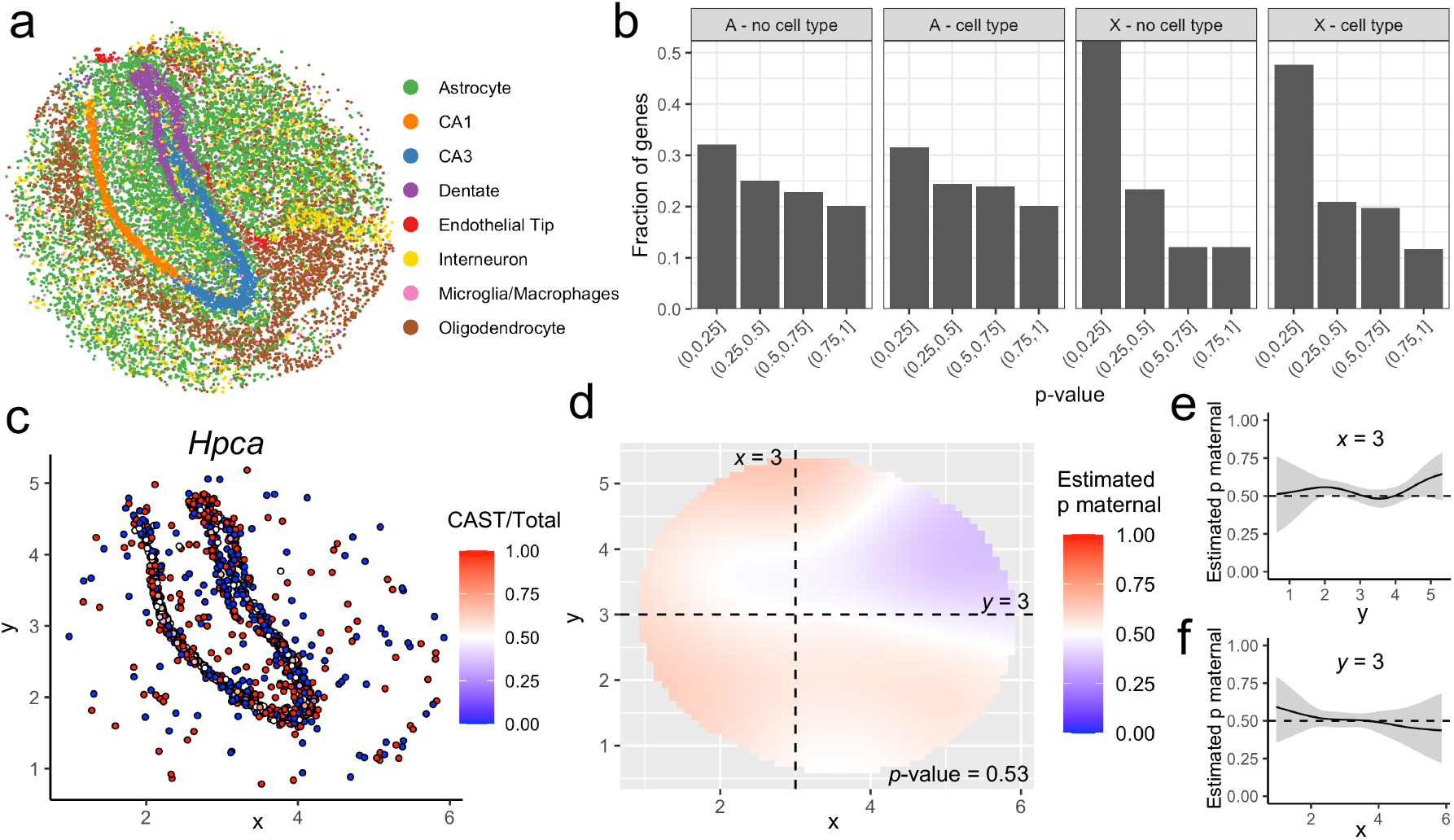
spASE identifies spatially significant ASE genes and smooths over sparse ASE spatial transcriptomics signal. **(a)** Map of cell types identified by RCTD in the Slide-seqV2 data generated in this study. Each point represents a pixel classified as a singlet of that cell type. **(b)** Distributions of *p*-values calculated by spASE for autosomal (A) and X-chromosome (X) genes in the real Slide-seqV2 data, not controlling for cell type (”no cell type”) and controlling for cell type (”cell type”). **(c)** Raw data for *Hpca*, showing higher coverage in the hippocampal formation and sparse coverage in the adjacent regions. **(d)** Estimated 2D maternal probability function for *Hpca*, with crosshairs indicating the *x* = 3 and *y* = 3 lines, along which point estimates and confidence intervals are plotted in **(e)** and **(f)**, respectively.

We then fit our model with and without the cell type covariates *X_ik_* in the model (see Methods). We found that, compared to autosomal genes, a higher proportion of X-chromosome genes had likelihood-ratio-test significance (Figure 2b), in concordance with the expected patterns of XCI in the X-chromosome. The *p*-value distribution for autosomal genes was closer to uniform distribution, indicative of less-frequent spatial ASE effects. After controlling for cell type, the autosomal distribution remained similar, while the distribution for the X-chromosome had a lesser skew, consistent with some the genes appearing to have spatial effects due to confounding with cell type effects.

Using a false discovery rate (FDR) threshold of *q* ≤ 0.01, we found ten genes with a spatially significant pattern, of which six were from the X-chromosome (Supplementary Table S1). However, after controlling for cell type, only three genes, two (*Tspan7* and *Plp1*) on the X chromosome and one (*Sst*) autosome, were signficant. Other genes, including *Nrip3* and *Ptgds*, were no longer significant after controlling for cell type, indicating that cell type differences were the main driver of spatial ASE for these genes.

spASE accounts for biological and technical noise to avoid detecting false positive ASE. For example, the gene *Hpca* was determined by spASE to not have signifcant spatial ASE (Figure 2c,d, *p*-value = 0.53). Although *Hpca* is highly expressed in the hippocampal formation, sparse expression in the adjacent regions resulted in noisier estimates and wide confidence intervals outside the hippocampus (Figure 2e,f). In general, such visualizations enabled by spASE allow for the assessment of both overall significance as well as position-specific significance across space.

### spASE detects spatial patterns of XCI across and within cell type in the mouse hippocampus

Next, we used spASE to estimate the maternal allele probability function for X-chromosome genes and found that the patterns for almost all significant X-chromosome genes were similar and anti-correlated with *Xist* expression (Figure 3a-c, Supplementary Figure S5), reflecting the expected mosaicism due to XCI by *Xist*. We also found that patterns of XCI were preserved within individual cell types. For example, *Tspan7*, which is relatively highly expressed in astrocytes (Figure 3d, S6), and *Plp1*, which is highly expressed in oligodendrocytes (Figure 3e, Supplementary Figure S7), were both estimated to have maternal probability functions anti-correlated with *Xist*.

**Figure 3:**
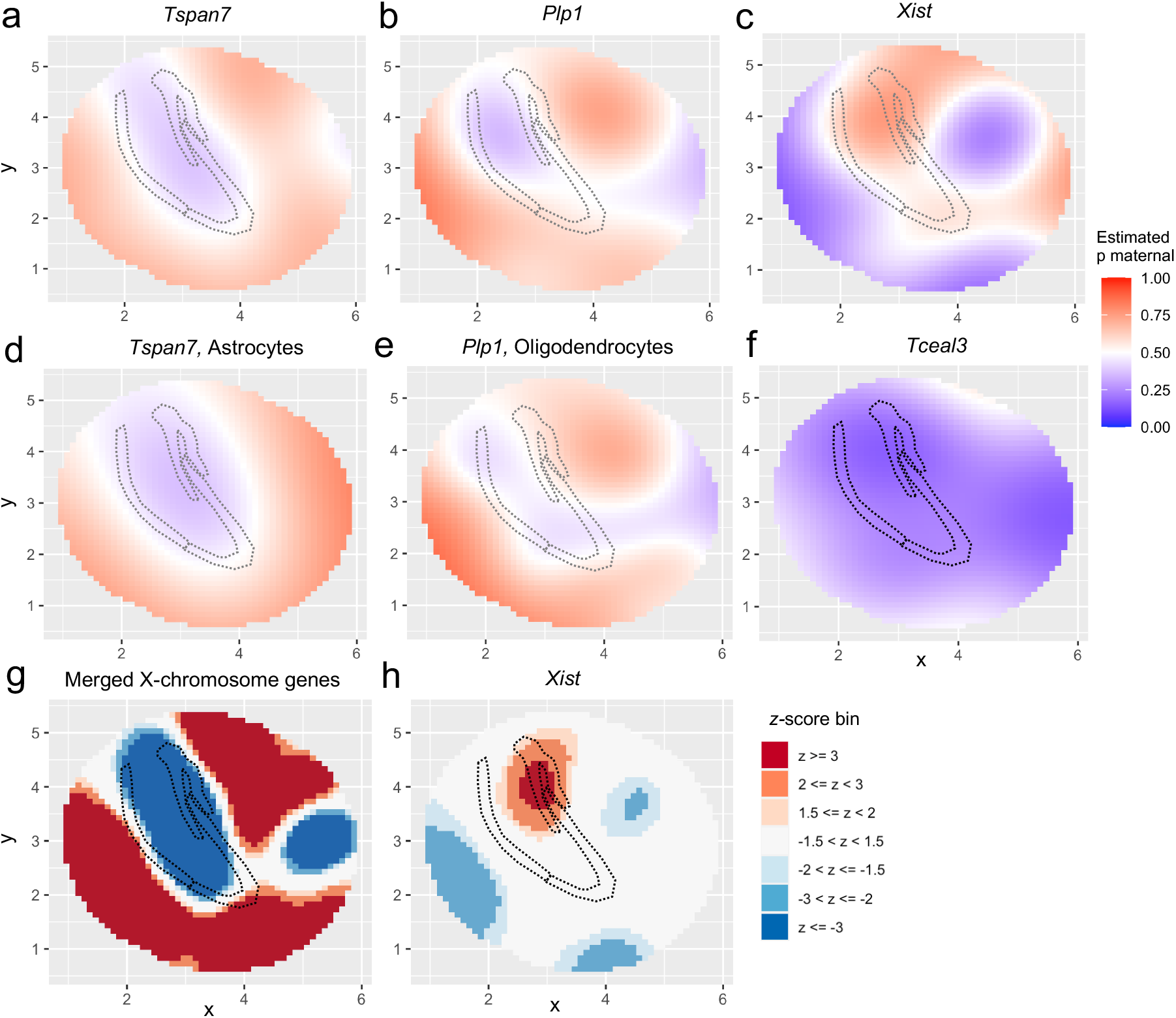
spASE detects spatial ASE in X-chromosome genes across and within cell type in the mouse hippocampus from Slide-seqV2 data. **(a)** Smoothed maternal allele probability functions for X-chromosome genes highly expressed in the mouse hippocampus and detected as significant (*q*-value ≤ 0.01): *Tspan7* and **(b)** *Plp1*. Red color indicates bias towards maternal, blue towards paternal, and white indicates both maternal and paternal alleles are present. The outline of the CA1, CA3, and dentate cell type regions is depicted in the dotted gray areas for reference. **(c)** Same as a-b for *Xist*. **(d)** Same as a-c but only using astrocyte pixels for *Tspan7*. **(e)** Same as a-c but only using oligodendrocyte pixels for *Plp1*. **(f)** Same as a-c for *Tceal3*. **(g)** 2D *z*-score plot computed from combining all *non-Xist* X-chromosome genes. **(g)** 2D *z*-score plot for *Xist*.

One X-chromosome gene, *Tceal3*, exhibited a strong paternal skew unlike the rest of the X-chromosome (Figure 3f, Supplementary Figure S8). However, the estimated *Tceal3* maternal probability still had a similar trend to the observed XCI pattern, with a high paternal bias around the hippocampus and a near-biallelic pattern in the periphery. We investigated the *Tceal3* locus and found that another nearby gene less than 100kb away, *Morf4l2*, also exhibited a strong paternal bias. *Tceal6*, a paralog of *Tceal3*, also showed a paternal bias in a similar pattern to that of *Tceal3*; however, other genes in the *Tceal* family, such as *Tceal5*, did not show the same bias (Supplementary Figure S8).

We then constructed a consensus XCI map by combining the UMI counts of all of the X-chromosome genes excluding *Xist* and fitting the model on the merged spatial profile. We visualized the significance at a region-specific level by computing and plotting *z*-scores in 2D (Figure 3g, Supplementary Figure S5). We found that the region significant for paternal X-chromosome expression was located precisely around the CA1, CA3, and dentate cell-type layers of the hippocampus as well as around a cluster of interneurons, and was anti-correlated with *Xist* (Figure 3h). *Xist* had fewer spatial regions reaching significance, reflecting its lower UMI coverage and thus wider confidence intervals in most areas (Supplementary Figure S9).

### spASE identifies cell type-driven spatial ASE in the autosome of the mouse hippocampus

We next investigated autosomal spatially significant genes with spatial ASE that was explained by cell type-driven ASE. Recall that several genes, including *Nrip3* and *Ptgds*, no longer possessed significant spatial ASE after controlling for cell type, indicating cell type-driven ASE. To quantify such cell type-driven ASE, we used spASE to estimate the overall maternal allele probability for each cell type, revealing several genes previously unknown to exhibit cell type-specific ASE (Figure 4a). For these genes, spASE’s estimated spatial ASE patterns were primarily driven by cell type localization distributions (Figure 4b-d). For example, *Nrip3*, one of the most statistically significant autosomal genes, had a high maternal bias in CA1, CA3, and dentate cell types, driving a strong maternal signal observed in the *z*-score plot (Figure 4b). *Ptgds*, which was highly expressed in both oligodendrocytes and endothelial tip cells, had a strong paternal bias in oligodendrocytes but not endothelial cells (Figure 4c,e). *Sst* exhibited a maternal bias which was enhanced in an interneuron subtype localizing primarily outside of the hippocampus (Figure 4d; Supplementary Figure S10).

**Figure 4:**
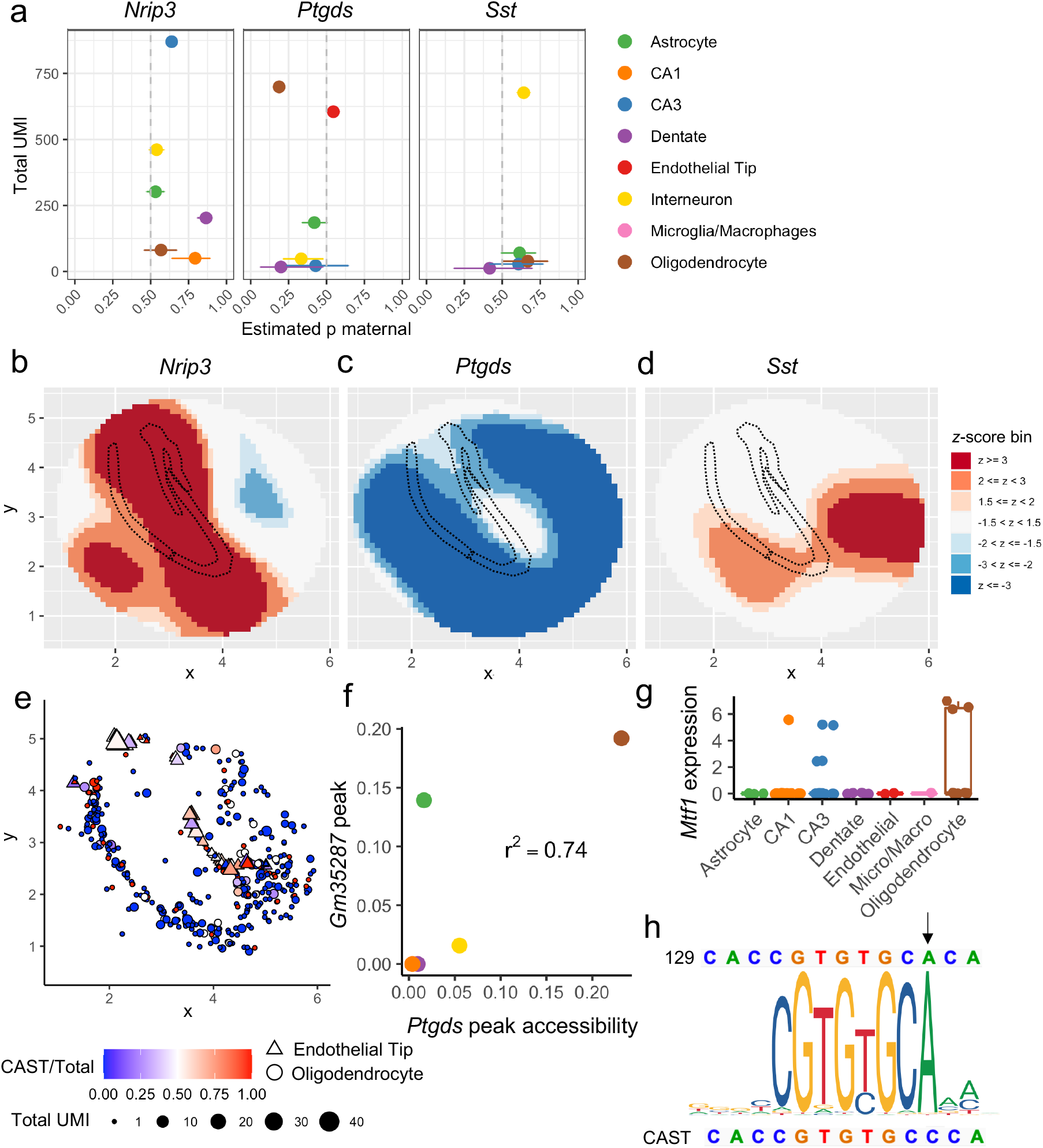
spASE identifies cell type-driven spatial ASE in the autosome. **(a)** MLEs and associated confidence intervals for the maternal probability *p* for three of the top autosomal gene hits (*q*-value ≤ 0.01), *Nrip3, Ptgds*, and Sst. x-axis: total UMI counts summed across all pixels. **(b)** 2D *z*-score plot for *Nrip3*. The hippocampal formation is outlined with dotted black lines. **(c-d)** Same as b for *Ptgds* and *Sst*, respectively. **(e)** Raw data for *Ptgds* for endothelial tip (triangles) and oligodendrocyte (circles) pixels. Size of the point indicates the total number of UMI present at that pixel, and color indications the fraction of the total UMI that were from the CAST (maternal) allele. **(f)** Average sci-ATAC-seq peak accessibility of the *Ptgds* promoter peak and the nearby (~8kb away) peak in *Gm35287* for the cell types overlapping between the sci-ATAC-seq data set and the Slide-seq data. **(g)** Single-cell RNA-seq expression for *Mtf1* from the Mouse Brain Atlas [43]. Each point represents a cluster that was classified as one cell type. **(h)** Position weight matrix for PB0044.1 (*Mtf1*) with 129 (paternal) and CAST (maternal) reference sequences shown on the top and bottom, respectively. Black arrow points to the SNP position of interest.

We examined whether this high cell type-specificity of ASE for these genes could be explained by genetic differences between the CAST (maternal) and 129 (paternal) mouse strains. Specifically, we investigated if a SNP could alter the binding affinity of a cell type-specific transcription factor at either the promoter or a putative enhancer. We analyzed single-cell ATAC-seq (scATAC) data from the mouse hippocampus [44] and searched for instances of SNPs overlapping known transcription factor binding site (TFBS) motifs in peaks within 50kb upstream and downstream of each gene (Methods). We found a peak in the promoter of *Gm35287* approximately 8kb upstream of *Ptgds* (Supplementary Figure S11), which is predicted to have a TSS-distal with enhancer-like signature for *Ptgds* [45]. Furthermore, we found that this peak has a high co-accessibility (*r*^2^ = 0.74) with the *Ptgds* promoter peak for cell types in common between the scATAC data and our spatial data (Figure 4f). In particular, both peaks are preferentially open in oligodendrocytes. We found a SNP, rs8255993, overlapping a known transcription factor motif, PB0044.1, which corresponds to the gene *Mtf1* (Supplementary Figure S11), which is also highly expressed in oligodendrocytes (Figure 4g) [43]. The SNP is A in the paternal strain and C in the maternal, and this position has a strong A signal in the TFBS position weight matrix for *Mtf1* (Figure 4h). Thus, the preferential binding of *Mtf1* to the paternal allele at this distal enhancer is a likely mechanism driving the paternal bias of *Ptgds* observed in oligodendrocytes.

## Discussion

Allele-resolved spatial transcriptomics suffers from high sparsity in comparison to bulk and single-cell sequencing, and confounding between cell type and spatial location present further statistical challenges. Here, we describe a statistical approach and software (spASE) which allows for estimating and visualizing 2D allele probabilities for sparsely expressed genes, as well as for testing spatial significance while controlling for user-specified covariates such as cell type. Through simulations, we demonstrate that our method maintains high power to detect ASE even with as few as 100 pixels and as low as 1 UMI per pixel for a given gene. We generated Slide-seqV2 data from an F1 female CASTx129 mouse hippocampus and show that our method recovers known patterns of XCI both within and across cell type (Figure 3). We further show that our method can identify cell type-specific ASE, which if not accounted for can be confused with spatial signal (Figure 4).

The primary *in situ* validation of our method was in the X-chromosome, where we found the same pattern of XCI both within and across cell types for multiple genes. XCI is thought to occur early in embryonic development in female organisms, before cell type differentiation [4], and the maternal and paternal chromosomes are thought to be equally likely to be inactivated. Thus, the pattern we observed in our data likely reflects randomly determined XCI in the early mouse embryo that propagated through to the adult hippocampus. This phenomenon can potentially explain why the X-chromosome p-value distribution was slightly but not fully affected by controlling for cell type (Figure 2b), as some nearby cell types may be derived from the same X-inactivated progenitor cell. Notably, *Tceal3* exhibited a strong paternal bias, but still had a spatial pattern that was similar to the general XCI pattern we observed in other X-chromosome genes (Supplementary Figures S5, S8). Another nearby gene, *Morf4l2*, also exhibited a paternal bias. Thus, the pattern we observed in *Tceal3* may be the combined result of XCI and another form of epigenetic imprinting.

One limitation of our spatial ASE analysis is that low UMI coverage limits the spatial resolution of ASE estimates. For example, within the XCI analysis, *Xist* was lowly expressed (Figure 3h); however, for genes with higher coverage, such as *Plp1*, it was possible to resolve the spatial ASE function further by increasing the degrees of freedom used to construct the 2D basis functions. Due to our limited spatial resolution, although we detected spatial patterns of XCI between the hippocampal formation (paternal bias) and surrounding areas (maternal bias), it is likely that increased statistical power would be achieved and higher-resolution spatial patterns would be uncovered given a higher-coverage dataset.

Similarly, although we found multiple instances of differential ASE across cell types as previously observed [20, 46], our analysis did not detect any spatial ASE in autosomal genes not explainable by cell type. We note that the statistical power was lower for the detection of spatial effects compared to the detection of cell-type differences. It is possible that autosomal spatial ASE effects might be detected given increased coverage and sample size.

We found that *Sst* exhibited a strong maternal bias for interneurons, particularly for a subtype located outside of the hippocampal formation with high *Sst* expression (Supplementary Figure S10). *Sst* is a well-known neuropeptide expressed throughout the brain which has been studied in the context of various neurological diseases [47]. However, we were not able to detect a likely cell type-specific transcription factor with a nearby binding site that was affected by strain-specific genetic variation as we did for *Ptgds*, although it is possible that the bias may only affect this subtype which is not represented in the scATAC-seq data set we used. Also, note that *Sst* exihbited low levels of expression in other cell types, which limited statistical power. Overall, these findings demonstrate that our method is broadly applicable for ASE discovery in spatial transcriptomics. Our rigorous computational approach will inform future analyses on the variability and biological mechanisms driving spatial and cell type-specific ASE.

## Methods

### Slide-seqV2 of CAST/EiJ x 129S1/SvImJ F1 mice

We obtained a female CAST/EiJ x 129S1/SvImJ (CASTx129) mouse from Jackson laboratories. The CASTx129 cross contains ~23 million SNPs, or approximately 1 SNP for every ~ 110 bp [48, 49]. This SNP density is approximately tenfold the SNP density in human cells and thus provides high resolution to interrogate ASE. Slide-seqV2 was performed as described previously [21, 22] on two adjacent, 10um-thick coronal slices of the hippocampus.

### Alignment of Slide-seqV2 data

We generated a pooled CASTx129 transcriptome using the command `create-hybrid` from the EMASE [50] software on the CAST and 129 transcript fasta files downloaded from ftp://churchill-lab.jax.org/software/g2gtools/mouse/R84-REL1505/. We then aligned 150bp reads to this pooled transcriptome with bowtie2 [51] using the parameters `-k 4 -p 16 --very-sensitive`. We used a custom script (https://github.com/lulizou/spASE/blob/master/scripts/processBowtie2.py) for processing the aligned BAM file [52] to create a gene UMI count matrix only from reads that uniquely aligned to one gene and one allele. We restricted attention to alignments with 3 or fewer mismatches and only considered alignments that had the fewest number of mismatches for that read. We overlaid data from the two slices by rotating and shifting the slices to overlap according to the location of the hippocampal formation.

### Beta-binomial model for allele-specific expression in spatial transcriptomics

Let *n_gi_* denote the observed total counts of gene *g* at cell or pixel *i* and *Y_gi_* denote the observed maternal allele UMI counts for gene *g* at cell or pixel *i*. Let λ*_gi_* denote the unknown mean probability of observing a maternal allele for each transcript of gene *g* at pixel *i*. We assume *Y_gi_*|λ*_gi_* ~ Binomial(*n_gi_*, λ*_gi_*), where *n_gi_* is observed and λ*_gi_* is a random variable, independently distributed (conditional on *ϕ_g_* and *p_gi_*, defined below) for each gene *g* and pixel *i*. We further assume that λ*_gi_* follows a beta distribution with mean *p_gi_* and variance *ϕ_g_p_gi_*(1 – *p_gi_*). The likelihood of this model for a single gene *g* can be written as

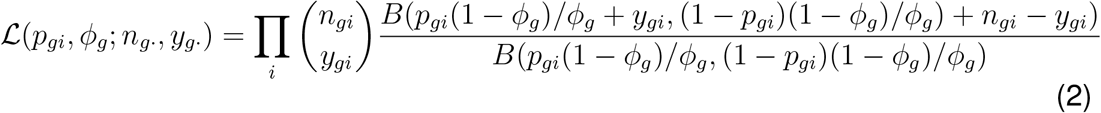

where *B* denotes the beta function. We used maximum likelihood estimation to obtain the estimates 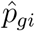 and 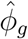 for each gene and the associated standard errors, determined from the Fisher information. When using this model for single-cell data to estimate ASE for a single gene, *p_gi_* is assumed to be the same for all cells *i*; when estimating *p_gi_* for spatial transcriptomics, *p_gi_* is assumed to be dependent on pixel *i* as described below. Assuming asymptotic normality, we used these point estimates and standard errors to construct 95% confidence intervals for 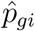. Finally, we used the Benjamini-Hochberg [53] procedure to produce *q*-values to control the false discovery rate.

In the spatial setting, for each gene *g*, we model *p_gi_* as a smooth spatially-varying function. Specifically, we used thin plate regression splines [36, 40, 41] to estimate smooth maternal allele probability surfaces for each gene. Thin plate regression splines allow estimation of a smooth function of 2D coordinates. The number of basis functions *d* determines the smoothness, with lower values of *d* corresponding to smoother functions. Choice of *d* depends on the sparsity of the data, since lower values of *d* reduce the variance, but at the risk of introducing bias. In the analysis presented here, we kept *d* constant across genes to ensure comparability. For our sparse Slide-seqV2 data, we found that *d* =10 provided enough complexity to model allelic patterns in the hippocampus sample examined here while also maintaining power to detect significant differences, and we used *d* =15 when plotting estimated maternal allele probability functions using all pixels. We also demonstrate reproducibility of results (i.e. genes detected as having a significant spatial pattern) across a range of values for *d* (Supplementary Tables S1–S4). In practice, we recommend visualization of the estimated probability function and confidence intervals (e.g. Figure 2c-f) to guide selection of *d*.

In the model for cell type-specific spatial ASE detection, the term *θ_gk_* corresponds to the effect size for cell type *k* for gene *g*. If no cell type annotations are available, or if a cell type effect does not exist (see likelihood ratio test below), then *θ_gk_* can be assumed to be the same for all *k*.

To test whether there was a significant spatial pattern beyond cell type, we assumed a baseline model with only cell types as covariates and performed a likelihood ratio test comparing model (1) to the baseline model, i.e. for each gene *g*, we compute

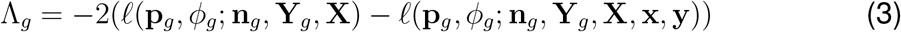

where *ℓ* is the log-likelihood computed from (1), **p**_*g*_ is the vector of maternal probabilities at each pixel *i* for gene *g*, **n**_*g*_ is the total number of UMIs at each pixel *i* for gene *g*, **Y**_g_ is the total number of maternal-derived UMIs at each pixel *i* for gene *g*, **X** is the *i* × *k* matrix of indicators of each cell type *k* at each pixel *i*, and **x**, **y** are the vectors of 2D spatial coordinates for each pixel *i*. This test statistic has an asymptotic 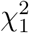 distribution, which we use to compute p-values for each gene.

For visualization, we plotted both the estimated smooth function for the MLE 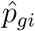 (Figure 3a-c) as well as 2D *z*-score plots (Figure 3d-e, Figure 4e). *Z*-score plots were calculated on an evenly spaced grid of points over the sample by taking the point estimate 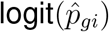 at each location and its associated standard error *s_gi_* and computing 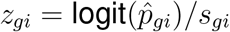.

### Computational implementation

spASE is implemented as an R package (https://github.com/lulizou/spase). We generated thin plate regression splines using the R package `mgcv` [40]. Specifically, we used the `smoothCon` function to construct spline basis functions. As basis functions can depend on the scale of the spatial covariates, we used normalized coordinates and also normalized the basis functions after construction by subtracting the mean and dividing by the standard deviation. We used the implementation of the beta-binomial likelihood from the R package `aod` [54].

We ran spASE in multiple modes: 1) not controlling for cell type, 2) controlling for cell type by allowing each cell type to have a different intercept term, and 3) allowing for each cell type to have a different spatial pattern. For 1), we experimented with using all pixels or only pixels confidently called as single cells (singlets). We found that using all pixels allowed us to increase our power and resolution for estimation of *p_gi_* and *ϕ_i_*, and for evaluating significance of spatial fits; thus, for visualization in our figures, we use all pixels, unless the figure is specifically denoted to be a single cell type. For results directly comparing to cell type models from 2) and 3), e.g. significant genes detected when controlling and not controlling for cell type, we use only pixels confidently classified in both cases to ensure comparable sample sizes of pixels.

### Simulation details

We simulated beta-binomial count data to evaluate the power, false positive rate, and *p*-values calculated using spASE. For each simulated gene, to construct random spatial ASE patterns, we used a random linear combination of basis functions calculated from the pixel locations of the Slide-seqV2 hippocampus data set using degrees of freedom *d* = 15. We first sampled a random number of total pixels *N* ∈ {50, 100, 250, 500} and used a fixed number of UMIs per pixel. The range of average UMIs per pixel for a single gene reached up to 14 in the real Slide-seqV2 hippocampus data set; for testing purposes, we used values ranging from 1 to 50. We drew the coefficients *θ_g_* of a random linear combination of basis functions from a standard normal distribution. Then, we chose a fixed overdispersion parameter *ϕ* ∈ {0.1, 0.3, 0.5, 0.8}, sampled the true binomial probabilities λ*_gi_* for each pixel location *i* from the beta distribution, and simulated counts from the binomial model.

To evaluate the asymptotic confidence interval coverage properties, we simulated ground truth data generated under the beta-binomial model without spatial covariates under a range of values for overdispersion, total coverage, and number of cells or pixels (Supplementary Figure S2). Specifically, we tested total UMI conditions based on three dataset-specific distributions for total UMI for a single gene across cells or pixels: Smart-seq3, which had generally high total UMI for each cell [31]; Slide-seq high, which had low counts with a high right skew; and Slide-seq low, which had mostly low counts (less than 5 per pixel, Supplementary Figure S3). We evaluated a range of values for the total number of cells or pixels the gene was captured on. We also tested a range of overdispersion values, namely *ϕ* ∈ {0, 0.1, …, 1}, and noticed the trend was the same as *ϕ* increased, thus we only show results here for *ϕ* = 0, 0.1, 0.8. We set the maternal probability at *p* = 0.5 and simulated 5,000 iterations for each condition.

### Prediction of cell types in spatial transcriptomics

We ran RCTD [23] to predict cell types in our Slide-seq data using a previously published single-cell RNA-seq reference for mouse hippocampus [42]. RCTD is a supervised approach that learns cell type profiles from a single cell reference and predicts cell type labels for spatial transcriptomics pixels using a Poisson log-normal model that accounts for platform effects between single-cell RNA-seq and Slide-seq data. We used a threshold of a likelihood difference of 100 between the minimum score and singlet score to classify singlets. After running RCTD, we filtered to pixels that were predicted to contain a single cell-type and added these cell type labels as the covariates in model (1) to perform all cell type-specific analyses.

### Mouse hippocampus scATAC-seq, cis-regulatory elements, and TFBS motifs

We analyzed previously published mouse hippocampus sci-ATAC-seq count matrices (GSE118987) [44]. The mice used in the study were the wild-type C57/B6 strain, and the data were aligned to the mm10 genome. We extracted the called peak annotations and counts using the command `scitools split`. To quantify accessibility of peaks, we computed the average count within cell types. We searched for known TFBS motifs within peaks with the command `matchMotifs` from the R package `motifmatchr` [55] using all motifs for *Mus musculus* in the JASPAR 2020 database [56]. We overlaid annotations of ENCODE cis-regulatory elements (cCREs) [45] for mm10 downloaded from the UCSC Table Browser [57] at the *Ptgds* locus and visualized the annotations using IGV [58].

### Mouse strain SNP data

We obtained gene-specific SNP annotations for 129S1/SvImJ and CAST/EiJ with respect to the mm10 reference using REL1505 of the Mouse Genomes Project (https://www.sanger.ac.uk/sanger/Mouse_SnpViewer/rel-1505)[59, 60]. For each gene of interest, we searched for SNPs overlapping the gene, within 50kb upstream of the gene start, and 50kb downstream of the gene end.

## Data availability

Slide-seqV2 data generated in this study will be made available upon publication.

## Acknowledgements

We thank members of the Irizarry lab for helpful discussions. LSZ was supported by an NSF Graduate Research Fellowship. FC acknowledges funding from NIH DP5OD024583, R33CA246455, NIH R01HG010647, the Burroughs Wellcome CASI fund, and the Searle Scholars award. RAI was supported by NIH grants R35GM131802 and R01HG005220.

## Author contributions

RAI, FC, and LSZ conceived of the study. TZ and EM generated the Slide-seqV2 data. LSZ, RAI, DMC, FC, TZ, and MJA analyzed the data. LSZ, RAI, DMC, TZ, and FC wrote the manuscript. All authors read and approved the final manuscript.

## Supplementary information

**Figure S1:**
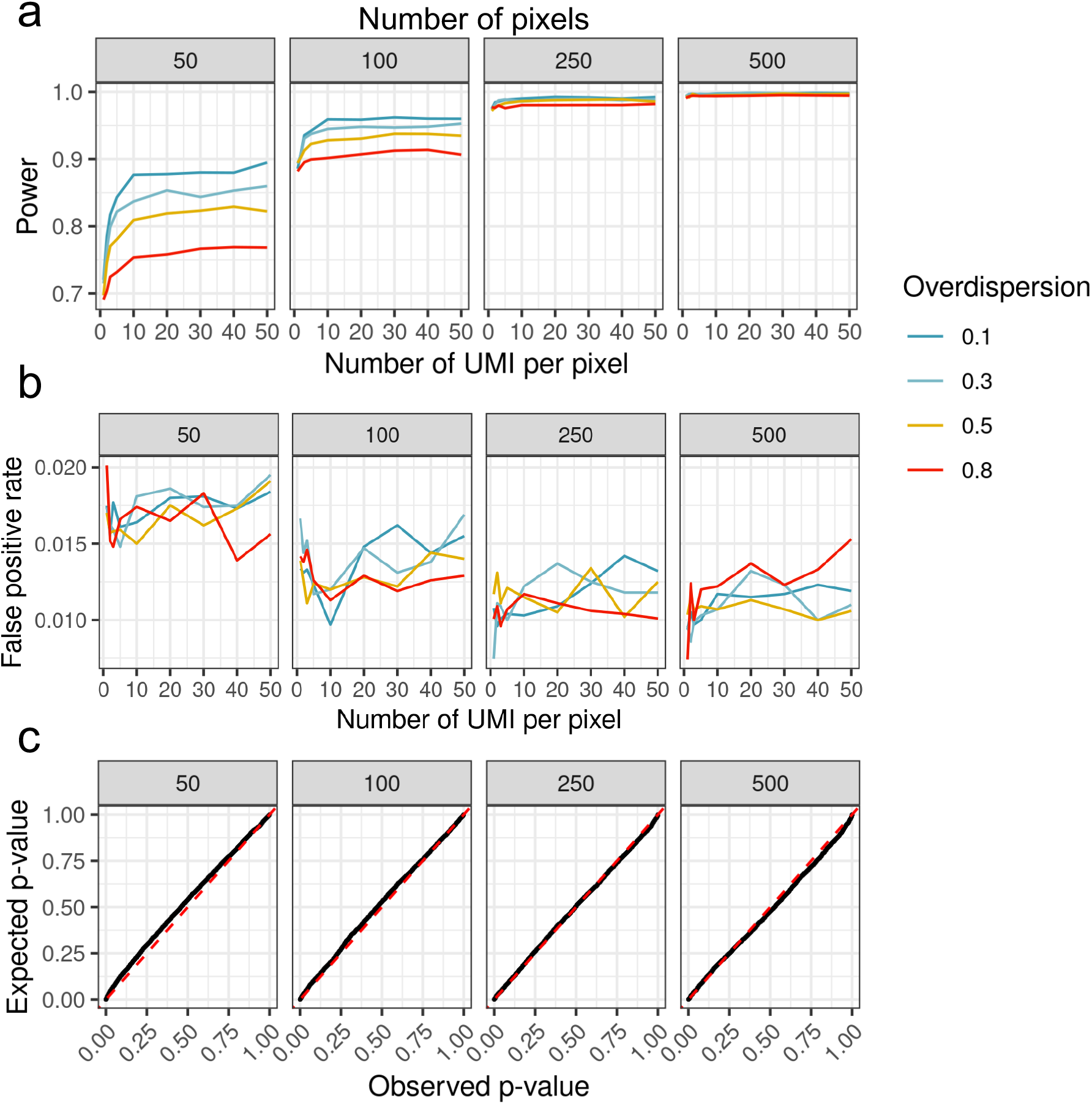
Spatial transcriptomic simulation results. **(a)** Power as function of number of pixels and number of UMI per pixel. x-axis is number of UMIs per pixel. Numbers in the gray panels indicate the number of total pixels. Curves are colored by the amount of overdispersion (*ϕ*) in the true model. **(b)** False positive rate as a function of number of pixels and number of UMI per pixel. **(c)** Expected *p*-values generated under a Uniform(0,1) distribution vs. observed *p*-values computed by spASE for the null case of no spatial ASE.

**Figure S2:**
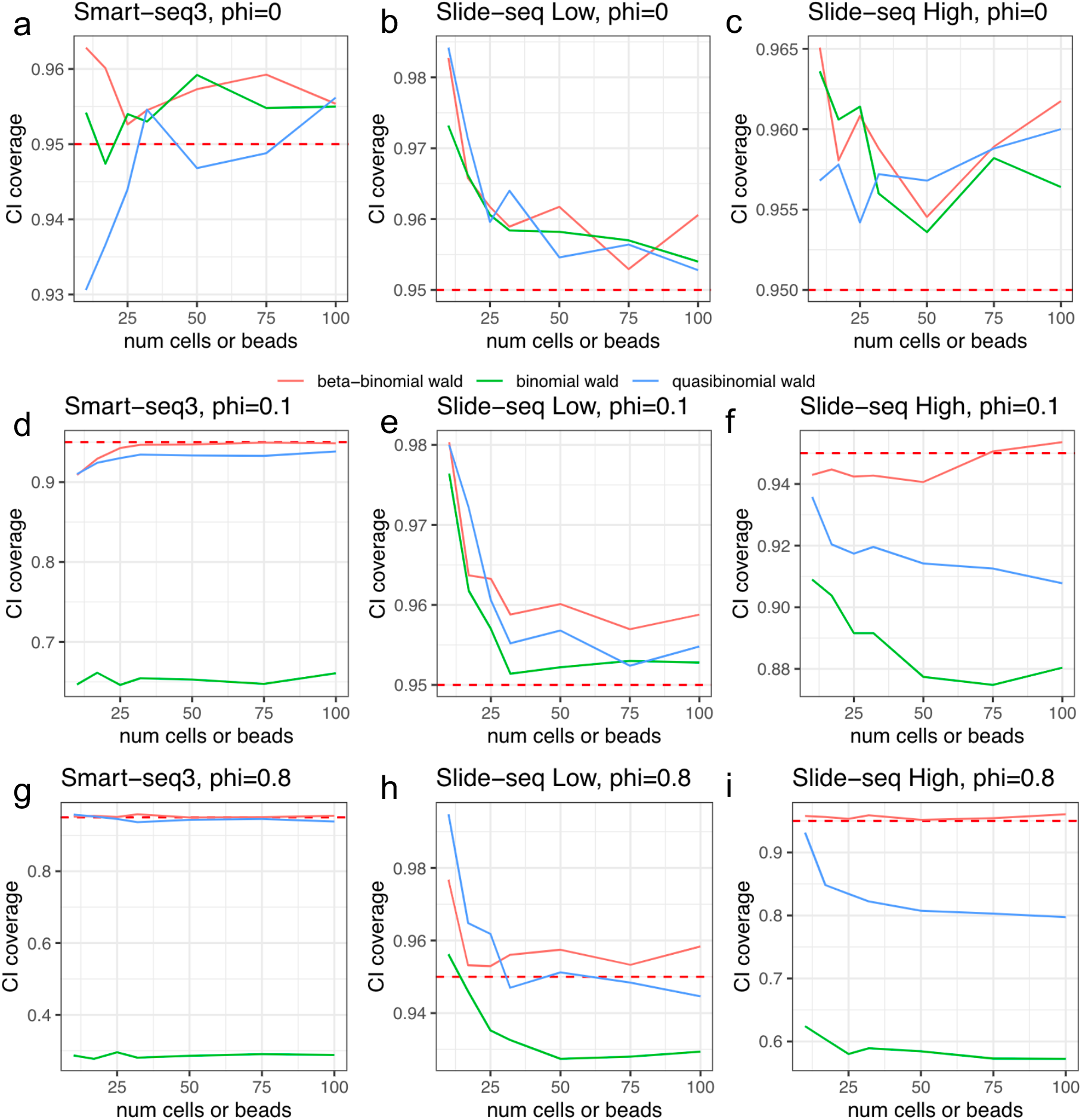
Simulation results for beta-binomial coverage probabilities as compared to the binomial and quasibinomial models. Data was generated from a beta-binomial model where each cell or bead had a total UMI count drawn from the total UMI count distribution from one of three settings, Smart-seq3 **(a,d,g)**, Slide-seq lowly expressed gene **(b,e,h)**, or Slide-seq highly expressed gene **(c,f,i)**. We also tested a range of values for overdispersion (phi): 0 **(a,b,c)**, 0.1 **(d,e,f)** and 0.8 **(g,h,i)**.

**Figure S3:**
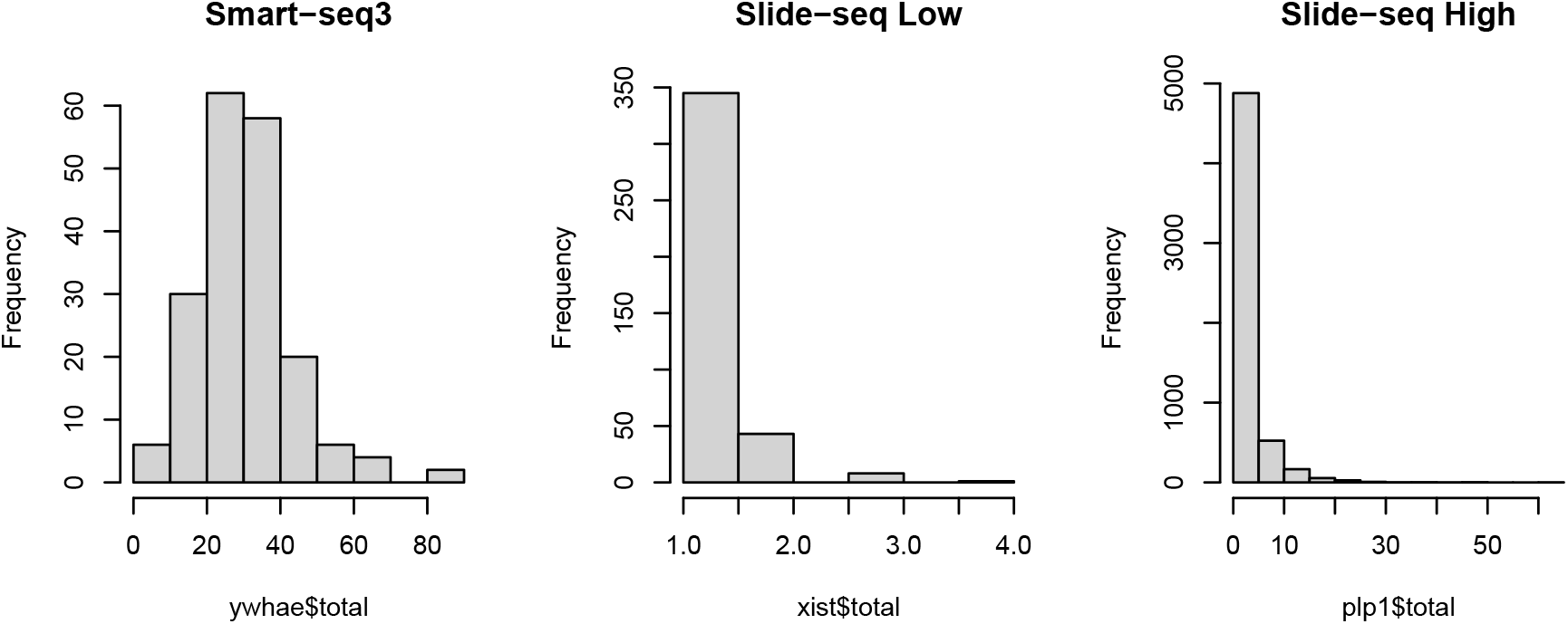
Total coverage distribution scenarios used for confidence interval coverage simulations shown in Figure S1. These were taken from genes to represent different sampling distributions for *n_gi_*, the total number of UMI per gene per cell (or pixel).

**Figure S4:**
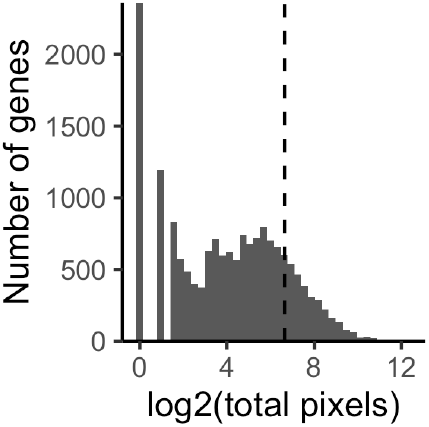
Histogram of total pixels that each gene had non-zero UMI counts for. The filtering threshold of 100 pixels per gene is shown with the dashed black line.

**Figure S5:**
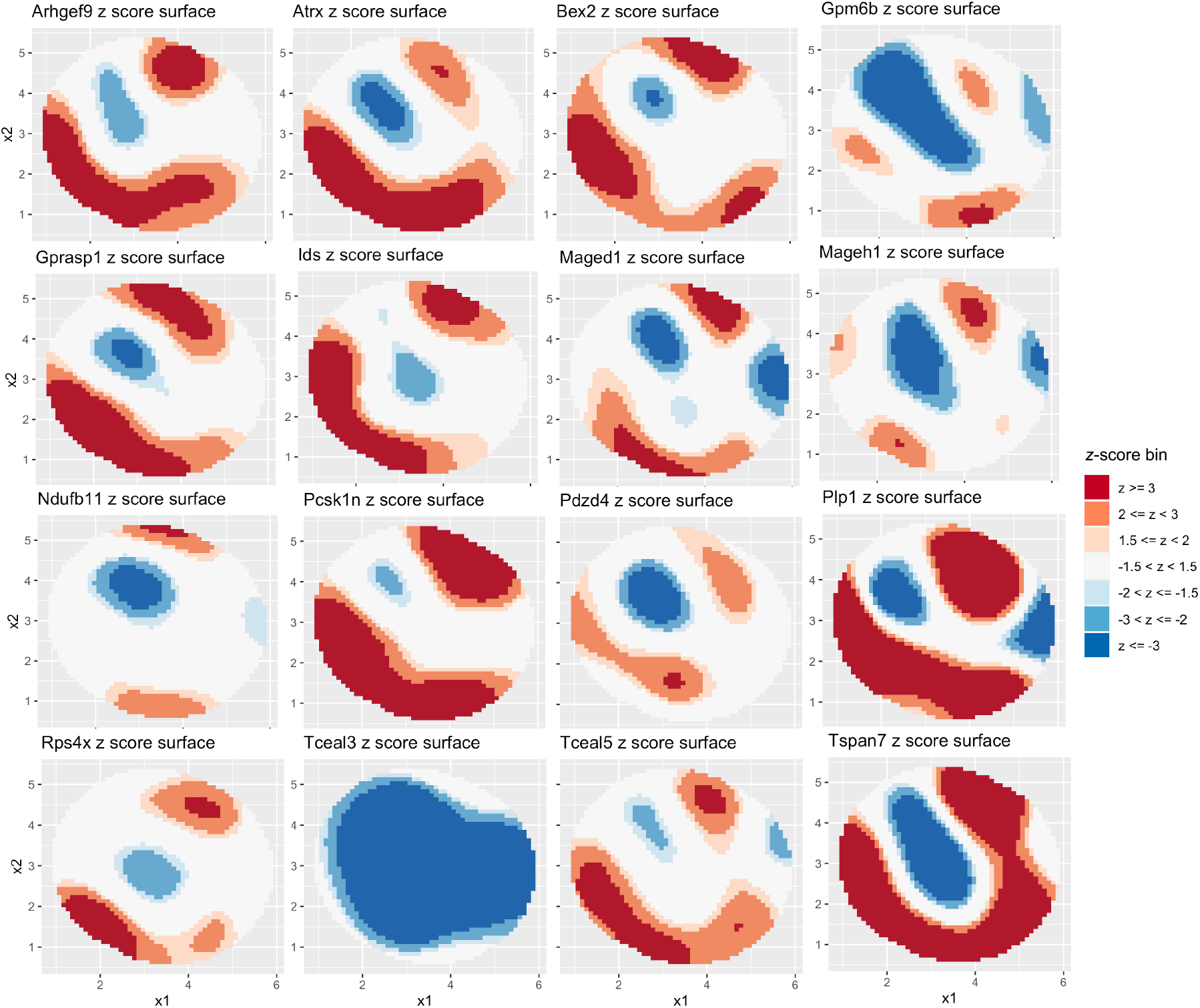
2D *z*-score plots for a sample of 16 highly expressed X-chromosome genes. Red color indicates bias towards the maternal (CAST) allele; blue indicates bias towards the paternal (129) allele.

**Figure S6:**
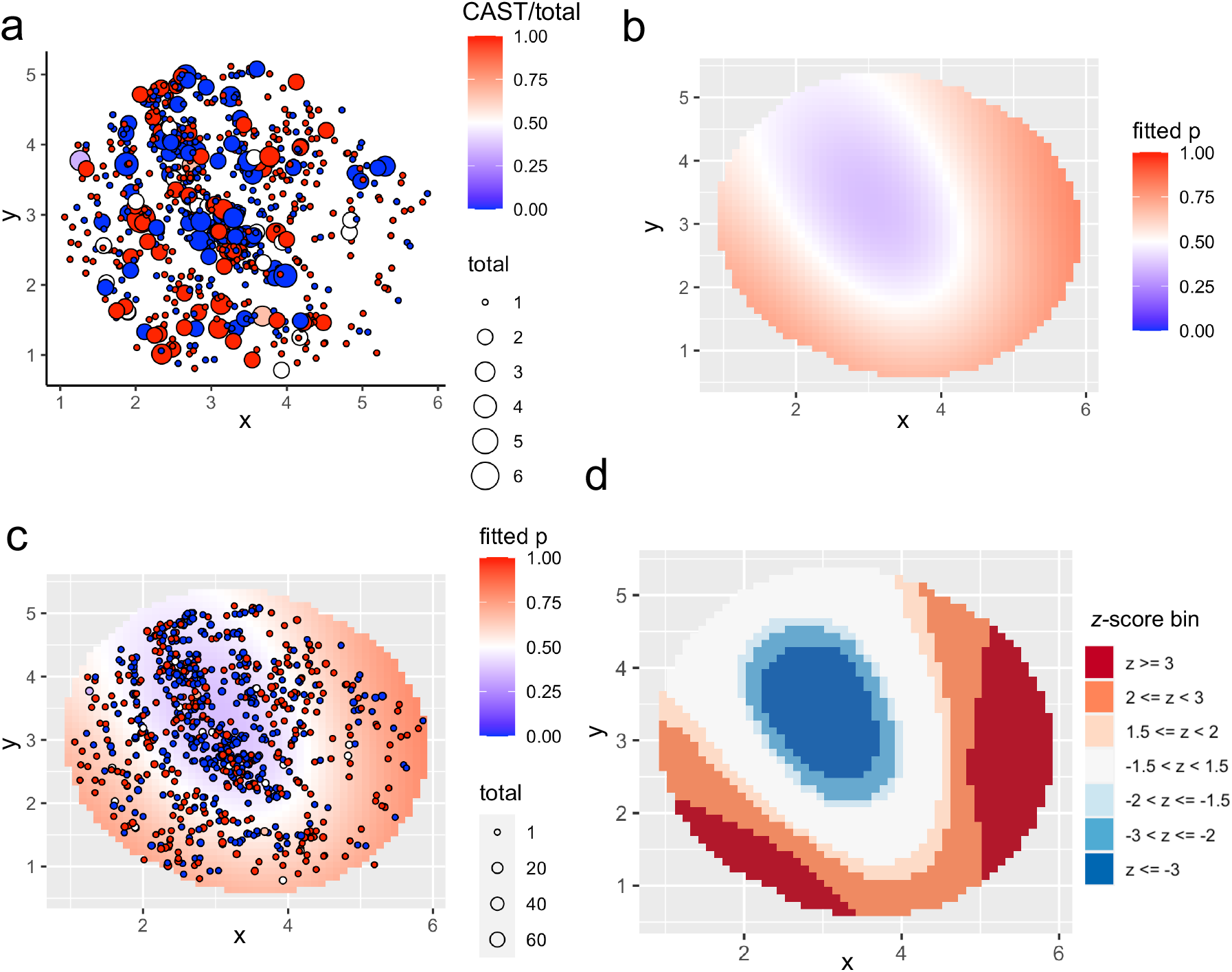
Within-astrocyte ASE for *Tspan7*. (**a**) Raw data for astrocyte singlets plotted using 2D coordinates for each pixel. The size of the point indicates the total UMI count for the gene *Tspan7* at that pixel. The color indicates the fraction of total UMIs that were from the maternal (CAST) allele. (**b**) Smoothed 2D maternal allele probability function (fitted p), estimated from the raw data shown in a using 5 degrees of freedom. (**c**) Overlay of data from a on the smoothed surface in b. (**d**) 2D *z*-score plot generated for the smoothed surface shown in b.

**Figure S7:**
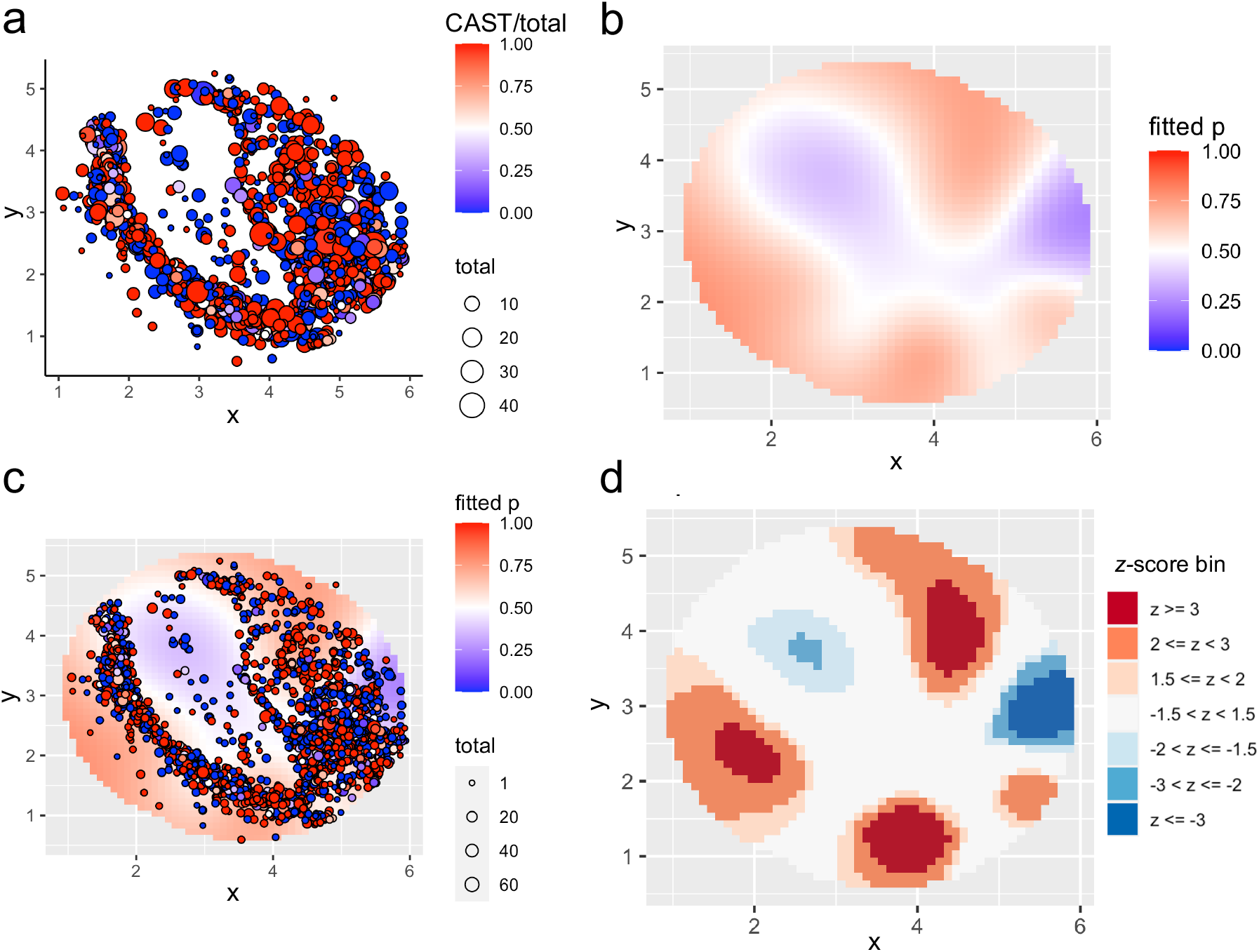
Within-oligodendrocyte ASE for *Plp1*. (**a**) Raw data plotted using 2D coordinates for each pixel. The size of the point indicates the total UMI count at that pixel. The color indicates the fraction of total UMIs that were from the maternal (CAST) allele. (**b**) Smoothed 2D maternal allele probability function (fitted p), estimated from the raw data shown in a. (**c**) Overlay of data from a on the smoothed surface in b. (**d**) 2D *z*-score plot generated for the smoothed surface shown in b.

**Figure S8:**
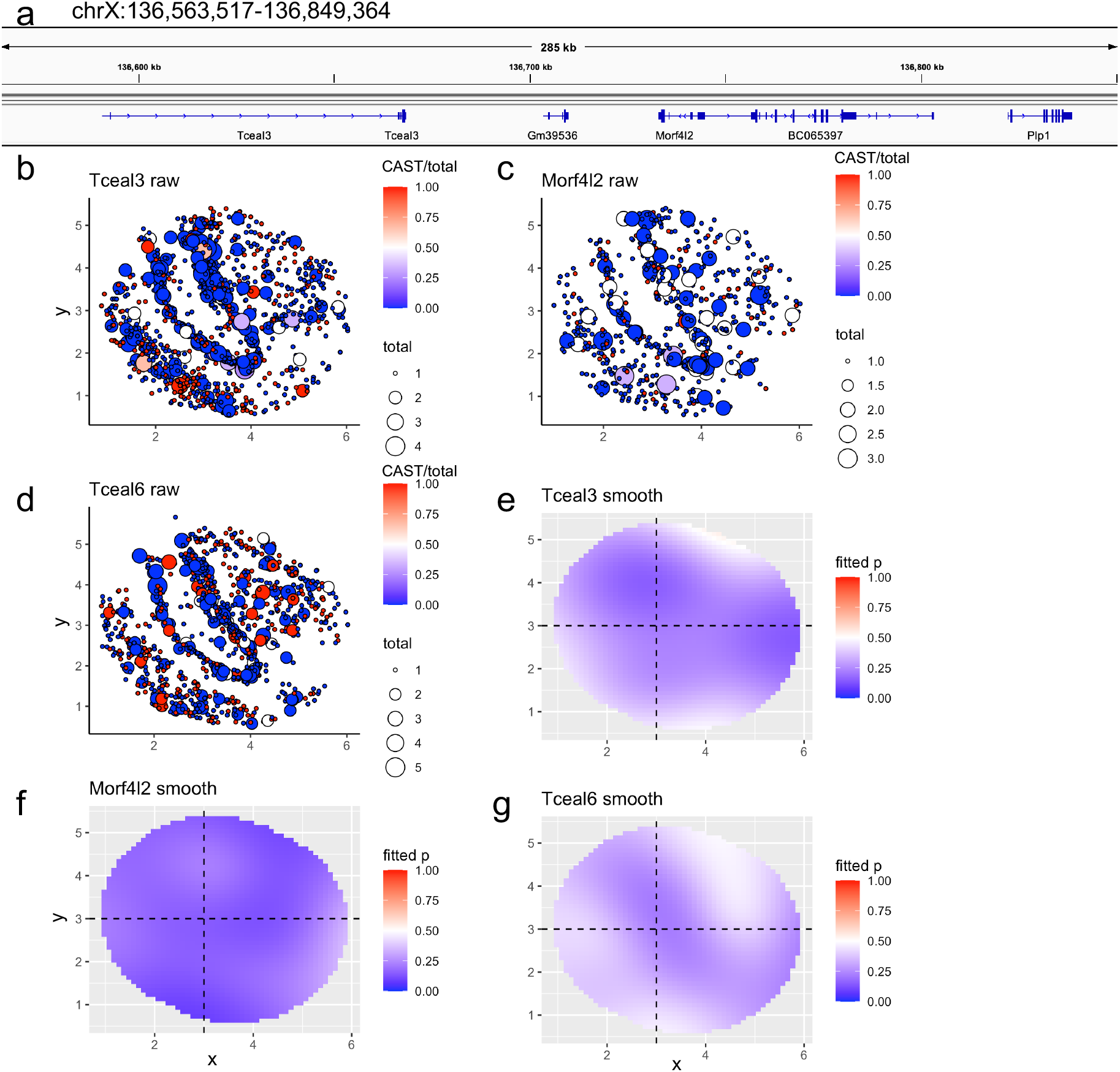
X-chromosome genes detected with a paternal bias. **(a)** IGV view of the *Tceal3* locus (coordinates are mm10). **(b)** Raw data for *Tceal3*, which was detected as having a significant spatial pattern (*q* ≤ 0.01). **(c-d)** Same as b for *Morf4l2* and *Tceal6*, which did not have a significant spatial pattern, but had paternal bias. **(e-g)** Smoothed maternal probability functions for *Tceal3*, *Morf4l2*, and *Tceal6*, respectively.

**Figure S9:**
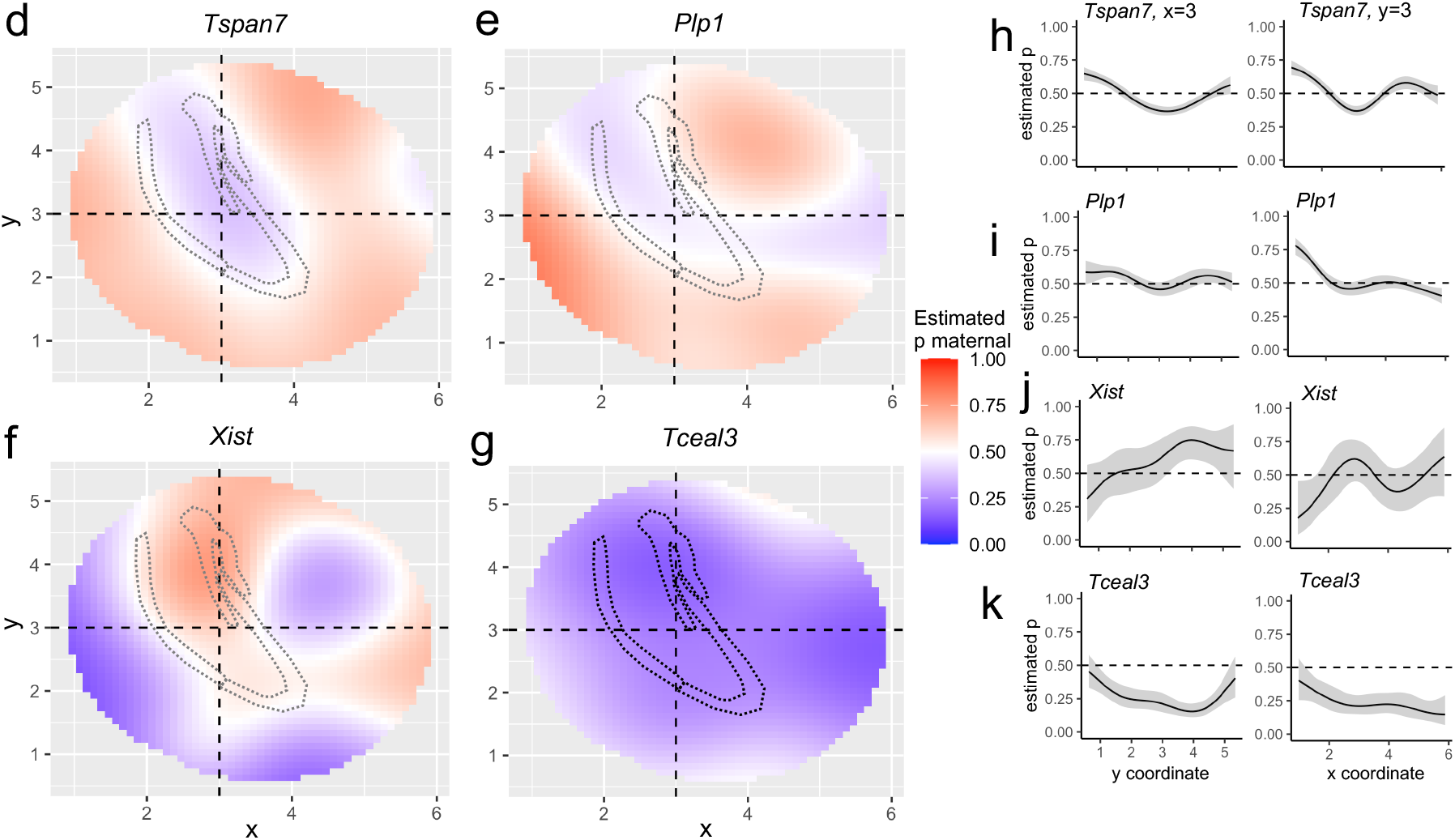
Confidence interval visualization for genes displayed in Figure 3. **(a-d)** Estimated maternal probability functions for *Tspan7, Plp1, Xist*, and *Tceal3* as shown in Figure 3. **(e-h)** Confidence interval visualizations in cross-sections along the *x* = 3 and *y* = 3 lines for each gene shown in a-d.

**Figure S10:**
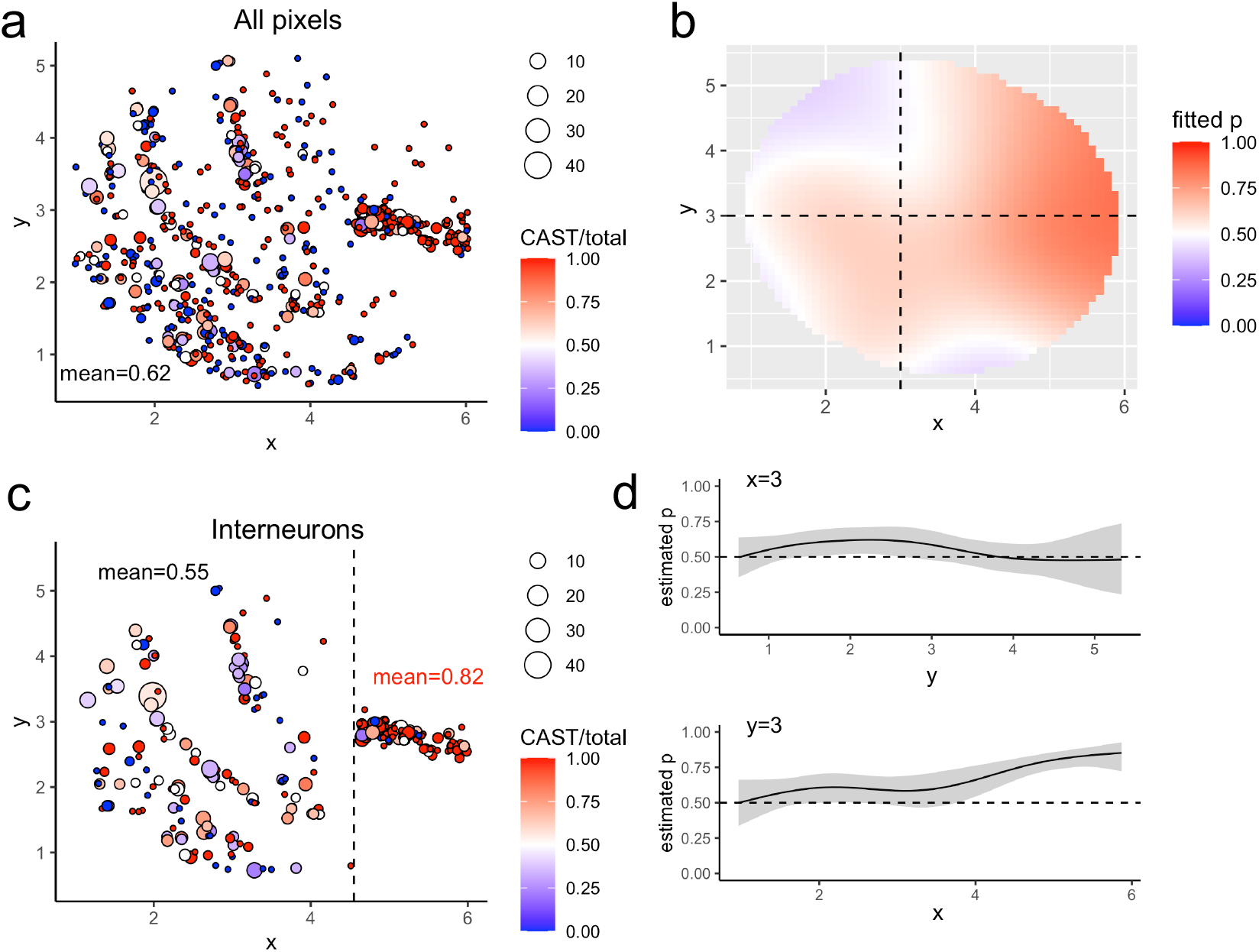
Interneuron ASE for *Sst*. (a) Raw data for each non-zero measurement pixel; color indicates fraction of total UMI that were maternal for each pixel, size of point indicates total UMI for that pixel. (b) Smoothed maternal allele probability surface. (c) Raw data for only interneuron singlets. Average expression for each boxed region is shown. (d) Confidence intervals from cross-hair slices in b.

**Figure S11:**
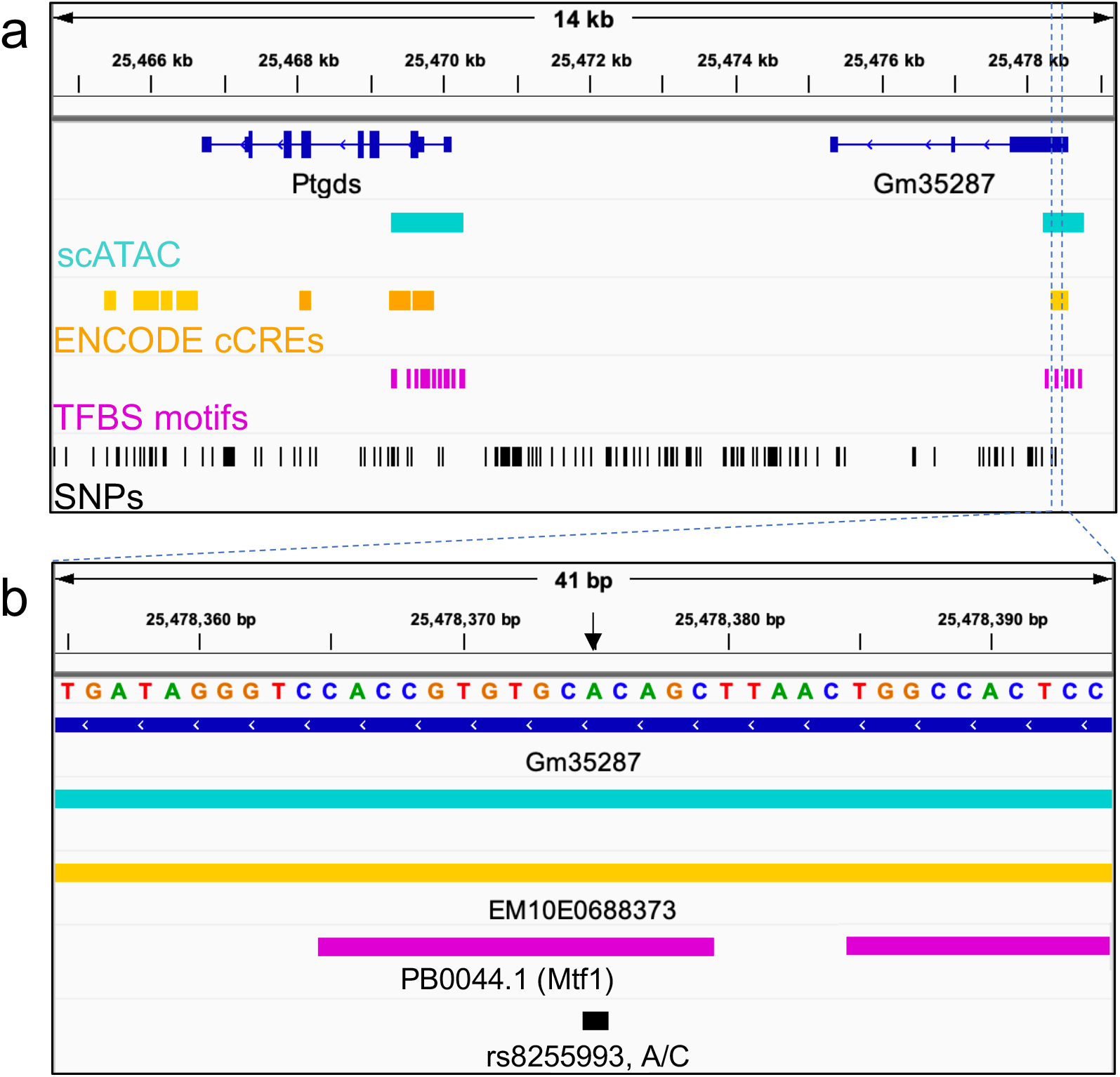
*Ptgds* locus with annotations. **(a)** IGV view (mm10) of the *Ptgds* locus showing the upstream *Gm35287* locus. Dark blue indicates Refseq gene annotation, cyan indicates peaks called from sci-ATAC-seq data from the mouse hippocampus, yellow denotes cis-regulatory elements (cCREs) from the ENCODE database (lighter yellow indicates distal-TSS enhancer-like signatures, darker yellow indicates proximal-TSS enhancer like signatures), magenta indicates predicted transcription factor binding site motifs within sci-ATAC-seq peaks, black indicates SNP locations for the CAST/EiJ and 129S1/SvmJ strains relative to mm10. **(b)** Zoomed-in genome browser view of the PB0044.1 motif (*Mtf1* gene) located in the peak overlapping *Gm35287*.

**Table S1:** Genes detected as spatially significant (*q*-value ≤ 0.01) in Slide-seqV2 of the mouse hippocampus, not controlling for cell type, degrees of freedom *d* = 10, restricting to pixels with a confident singlet classification by RCTD.

| | Gene | Total UMI | $\chi^2$ $p$-value | $q$-value | X-chr |
| --- | --- | --- | --- | --- | --- |
| 1 | Ptgds | 1584 | 0.00e+00 | 0.00e+00 | FALSE |
| 2 | Tspan7 | 4744 | 0.00e+00 | 0.00e+00 | TRUE |
| 3 | Plp1 | 12850 | 3.33e-16 | 4.53e-13 | TRUE |
| 4 | Nrip3 | 1963 | 8.39e-11 | 8.56e-08 | FALSE |
| 5 | Sst | 837 | 2.45e-07 | 2.00e-04 | FALSE |
| 6 | Pcsk1n | 1523 | 3.83e-07 | 2.60e-04 | TRUE |
| 7 | Rgs4 | 679 | 4.02e-06 | 2.34e-03 | FALSE |
| 8 | Atrx | 1096 | 5.37e-06 | 2.74e-03 | TRUE |
| 9 | Mageh1 | 362 | 9.04e-06 | 4.10e-03 | TRUE |
| 10 | Gpm6b | 1601 | 2.07e-05 | 8.44e-03 | TRUE |

**Table S2:** Genes detected as spatially significant (*q*-value ≤ 0.01) in Slide-seqV2 of the mouse hippocampus, not controlling for cell type, *d* = 5, restricting to pixels with a confident singlet classification by RCTD.

| | Gene | Total UMI | $\chi^2$ $p$-value | $q$-value | X-chr |
| --- | --- | --- | --- | --- | --- |
| 1 | Tspan7 | 4744 | 8.30e-14 | 3.41e-10 | TRUE |
| 2 | Nrip3 | 1963 | 4.82e-11 | 6.60e-08 | FALSE |
| 3 | Ptgds | 1584 | 4.18e-11 | 6.60e-08 | FALSE |
| 4 | Sst | 837 | 4.19e-09 | 4.31e-06 | FALSE |
| 5 | Rgs4 | 679 | 6.43e-07 | 5.29e-04 | FALSE |
| 6 | Lypd1 | 241 | 1.40e-05 | 9.61e-03 | FALSE |

**Table S3:** Genes detected as spatially significant (*q*-value ≤ 0.01) in Slide-seqV2 of the mouse hippocampus, not controlling for cell type, *d* = 15.

| | Gene | Total UMI | $\chi^2$ $p\text{-value}$ | $q\text{-value}$ | X-chr |
| --- | --- | --- | --- | --- | --- |
| 1 | Plp1 | 12850 | 0.00e+00 | 0.00e+00 | TRUE |
| 2 | Ptgds | 1584 | 0.00e+00 | 0.00e+00 | FALSE |
| 3 | Tspan7 | 4744 | 0.00e+00 | 0.00e+00 | TRUE |
| 4 | Nrip3 | 1963 | 4.89e-11 | 4.92e-08 | FALSE |
| 5 | Gstm7 | 326 | 1.51e-06 | 1.14e-03 | FALSE |
| 6 | Pcsk1n | 1523 | 1.70e-06 | 1.14e-03 | TRUE |
| 7 | Sst | 837 | 3.05e-06 | 1.76e-03 | FALSE |
| 8 | Gpm6b | 1601 | 4.94e-06 | 2.49e-03 | TRUE |

**Table S4:** Genes detected as spatially significant (*q*-value ≤ 0.01) in Slide-seqV2 of the mouse hippocampus, not controlling for cell type, *d* = 20.

| | Gene | Total UMI | $\chi^2$ $p\text{-value}$ | $q\text{-value}$ | X-chr |
| --- | --- | --- | --- | --- | --- |
| 1 | Ptgds | 1584 | 0.00e+00 | 0.00e+00 | FALSE |
| 2 | Tspan7 | 4744 | 0.00e+00 | 0.00e+00 | TRUE |
| 3 | Nrip3 | 1963 | 1.38e-10 | 1.80e-07 | FALSE |
| 4 | Gpm6b | 1601 | 1.08e-07 | 9.42e-05 | TRUE |
| 5 | Pcsk1n | 1523 | 1.21e-07 | 9.42e-05 | TRUE |
| 6 | Gstm7 | 326 | 4.06e-06 | 2.63e-03 | FALSE |
| 7 | Magt1 | 127 | 8.99e-06 | 5.00e-03 | TRUE |
| 8 | H1f2 | 105 | 1.66e-05 | 8.10e-03 | FALSE |

